# The Effective Inertia of the Lower Limb During Locomotion

**DOI:** 10.1101/2025.10.03.680415

**Authors:** Guanxiong Chen, Bastian Wandt, Helge Rhodin, Dinesh K. Pai

**Affiliations:** University of British Columbia; Linköping University; Bielefeld University

**Keywords:** Inertia Estimation, Musculoskeletal Modelling, Motion Reconstruction, Ground Reaction Force

## Abstract

Accurate modelling of inertial properties of human lower limbs is of great interest to many tasks, from gait analysis in biomechanics to motion tracking and control in computer animation. Previous work typically simplified the human musculoskeletal structure as a chain of rigid capsules, with muscle mass lumped with body segments. Such simplifications lead to errors in the inertia matrix, and the error propagates to torque and pose estimates. In this study, we developed a data-driven model to represent the joint-space inertia of the lower-body of a human in motion. The model does not make any assumptions other than that the estimated inertia matrices must be symmetric and positive definite. We show that a joint-space inertia matrix, estimated from synchronized motion and ground force data following foot strikes, reveals inertial coupling, and that estimated inertia matrices are bilaterally symmetric and motion-type dependent. These are properties which a rigid, mass-lumped inertia matrix fails to entail. Moreover, we show that the data-driven model fits to data better than the articulated rigid body inertia model, and that when used for reconstructing lower body kinematics estimated inertia yields more accurate and stable motion.

## Introduction

When a foot strikes the ground during locomotion, large contact forces and accelerations come into play. These are fundamentally linked to the effective inertia of the body, particularly the lower limb. However, the limb’s inertia changes in complex ways throughout locomotion. It is well known that joint articulation contributes to pose-dependent changes in inertia. Efficient methods for computing the pose-dependent inertia of articulated chains of rigid bodies, such as the Articulated Rigid Body (ARB) Model have been developed and widely used in graphics, robotics and biomechanics (Xiang et al., 2011; Shourijeh et al., 2017; Dao, 2019; Zell et al., 2020; Lam and Vujaklija, 2021; Ren et al., 2007; Sharif Razavian et al., 2019; Song et al., 2021; Clancy et al., 2023; Nakada et al., 2018; Jiang et al., 2019; Lee et al., 2019; Park et al., 2022).

Recently, however, it has been recognized that the ARB model is a poor approximation of how mass in a limb actually moves. Muscle mass is often connected, via light and stiff tendons, to distal limb segments, and therefore contributes to the effective inertia of those segments (Pai, 2010). A further complication is that neural activation of a muscle can affect how strongly its mass is coupled to the distal insertion, potentially causing the effective inertia to change with activity (e.g., walking vs. running). Accounting for the inertial contribution of muscles is challenging and can produce surprising effects (e.g., Verheul et al. (2023)).

Similar to prior work (Pataky et al., 2003; Venture et al., 2008; Damavandi et al., 2009; Chen et al., 2011; Hansen et al., 2014; Jovic et al., 2016), from motion capture and force plate data collected from 22 subjects during walking and running, we directly estimate each subject’s effective inertia under each motion. We focus on the effective inertia during the foot strike, or initial contact phase of the gait cycle, and use a data-driven approach to estimate a “correction inertia” not accounted for by the classical ARB model. We introduce methods to ensure that the estimated inertia is symmetric and positive-definite.

Our results show that the effective inertia at distal joints is greater than predicted by classical ARB models, and that inertia during running is significantly higher than during walking. These findings are consistent with the contribution of active muscle inertia to distal joints and highlight the importance of properly accounting for muscle mass in musculoskeletal models.

We implemented our methods in Python and MATLAB; The dataset we used is publicly available.

## Methods

### A. Correction Inertia

Prior work in human motion reconstruction and analysis (Zell et al., 2020; Zhang et al., 2024) typically formulate the dynamic equation of motion of a human kinematic model in generalized coordinates as

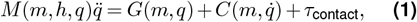

where *m* is the body mass of a human subject, *h* is the body height of the subject *M* (*q*) is the *D* × *D* (where *D* is the degree of freedom of the kinematic model) subject and pose-dependent joint-space inertia matrix of the subject, *q* is pose, *G*(*m, q*) is the pose and body mass-dependent gravitational force (GRF), 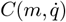 is the Coriolis force, *τ*_contact_ is ground reaction force and moment. *M* (*q*) is often habitually constructed from the articulated body (ARB) assumptions, but an analytically constructed ARB inertia cannot capture the complex geometry and mass distribution of body segments. Moreover, due to lumping of muscle mass to skeletal mass, the inertia of a joint would only be influenced by its distal segments, and the mass of a muscle will not contribute to the inertia of its distal joints, even if it should (Pai, 2010). To address the two problems, Wang et al. (2022); Verheul et al. (2022) levelled up details in analytical construction by approximating a muscle as a thread of mass points, with mass uniformly distributed among the points. We take a hybrid data-driven and analytical approach unlike the prior work: given measured body mass *m* and pose *q*, first of all we compute a joint-space inertia matrix under the ARB assumptions; then we use joint velocity and GRF to estimate a *correction inertia* matrix *M*_correction_(*q*) that accounts for factors that the ARB assumptions are unable to capture, including complex geometry, non-uniform mass distribution and inertial coupling

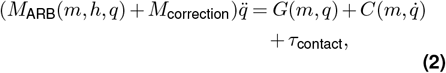

We refer to the element-wise sum of an ARB inertia matrix and the corrective inertia matrix as the joint-space *effective inertia* matrix. Our analysis will focus on properties of the effective inertia matrix and distinctions from the ARB inertia matrix.

### B. Dataset

We used the publicly-available human motion dataset from Zell et al. (2020). It contains 318 walking clips and 128 running clips collected from 22 subjects, with different gender and BMI’s; each clip consists of joint poses sampled via a Vicon Mocap System and contact forces sampled via AMTI force plates (Zell et al., 2020) at 10 ms intervals, from the first half of a gait cycle. To learn if inertia is motion-type dependent, we divided the dataset into the Walking Set and the Running Set, and estimated inertia on the two sets separately. Since the inertia matrix also depends on body mass proportion and pose, we further partitioned each of the two sets on a per-subject, per-pose basis: split into subsets, each made up of gait cycles from one subject, then divided the subset into left-foot and right-foot contact set. See Fig. 1 (a). We exclusively use data from *t*_0_ ≤ *t < t*_*L*_ = 130ms in a gait clip for estimation: see Fig. 1 (b). This is a period of time during which the ipsilateral foot is in contact with the ground and the contralateral foot has just lifted off, and it lasts for roughly 13% of a gait cycle (Bonnefoy-Mazure and Armand, 2015). Using data immediately following foot strike allows us to ignore velocity-dependent forces (e.g. the Coriolis force) and muscle control without incurring significant errors, because ground reaction force is sufficiently large to be dominant. Fig. 1 (c) shows the kinematic model used in Zell et al. (2020).

**Figure 1:**
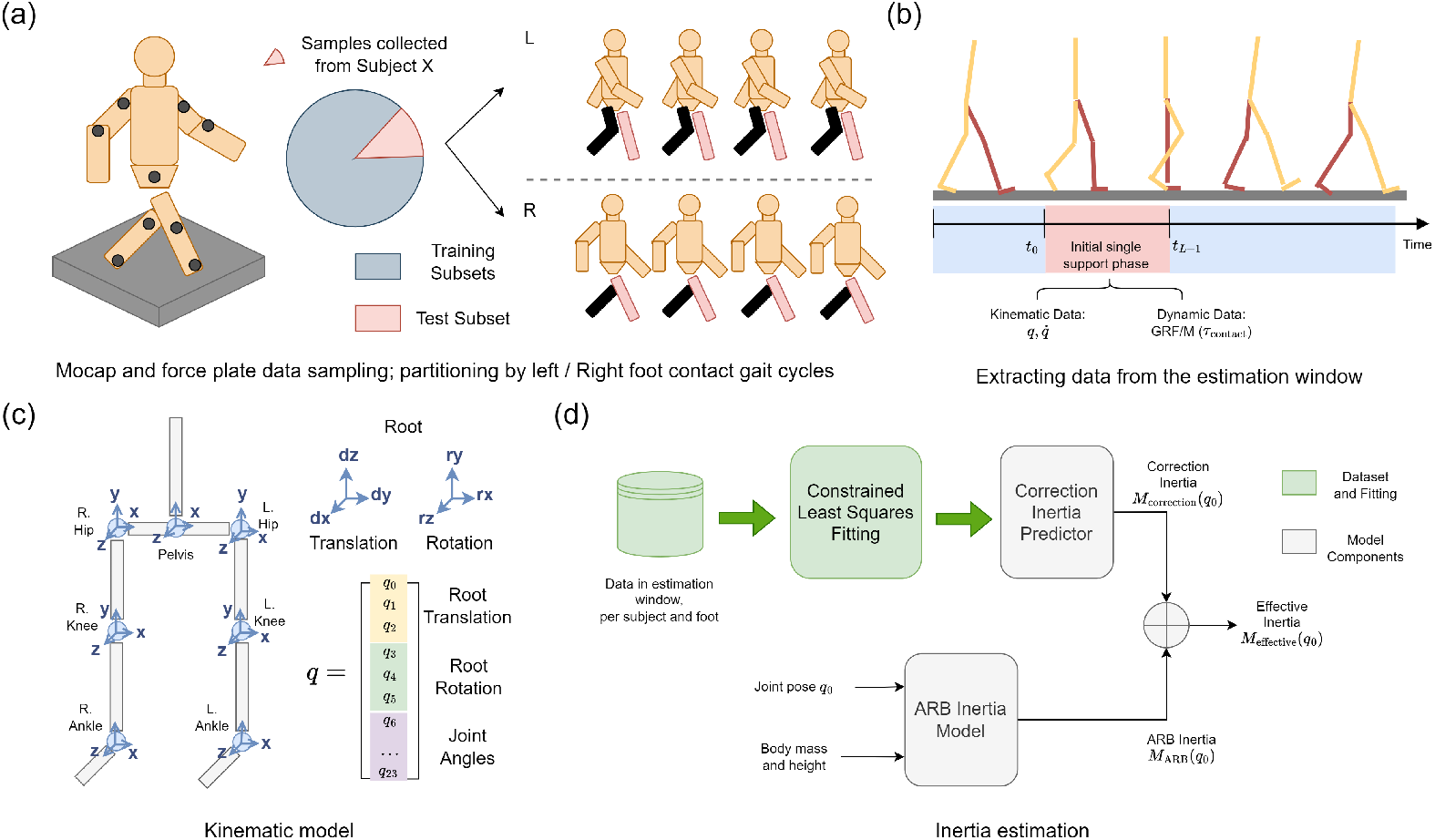
Method overview. (a) Curation of mocap and force plate data and the dataset partitioning scheme. (b) Extraction of data to use for inertia estimation. We take data from the first 130 ms of a walking or running gait after foot strike. (c) Kinematic model of the dataset and generalized coordinates to characterize the system. It consists of 24 degrees of freedom, including 6 degrees representing root translation and rotation, and 18 lower-body joint angles. We focus on analyzing inertial properties and reconstructing motion of lower body segments and the torso. (d) Components of the data-driven inertia model. Note that out of the 22 subjects from the dataset (Zell et al., 2020), only 13 of them have both walking and running gaits collected. Therefore, for experiments in which we need to compare walking and running inertia estimates we use only contact sets of the 13 subjects.

### C. The Data-driven Inertia Model

As illustrated in Fig. 1 (d), we can estimate a single effective inertia matrix for a particular subject-pose in a 130 ms estimation window. Starting from dsicretizing Eq. 2, for a *d* = 24 DoF human model at any timestep *t* = *t*_*i*_ in the window during *t*_0_ ≤ *t*_*i*_ ≤ *t*_*L*−1_, on velocity level

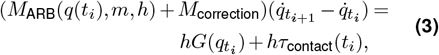

where *q* is the *D*-dimensional generalized coordinates, *G* is gravitational torque and *h* is the length of a timestep. We drop the Coriolis term since it is small relative to other terms during the ground contact phase. *M*_ARB_ is the ARB inertia matrix which we can construct from pose *q*(*t*_*i*_), body mass and height (see Sec. D); *M*_correction_ is the correction inertia matrix that is estimated specifically on pose *q*(*t*_*i*_) and this subject. We expect the effective matrix to be symmetric and positive definite (SPD), since a non-SPD effective inertia implies a non-positive eigenvalue which is physically implausible. So for each subset of multiple gait clips from the same subject and stance leg, we can divide it into a training set of *N* gait clips, a validation and a test set, and use the *N* training clips to fit a simple yet effective statistical model to estimate a correction inertia, by minimizing the difference between predicted and ground-truth per-step squared impulse *f* (*M*_correction_) : ℝ^*D*×*D*^→ℝ, using constrained least squares

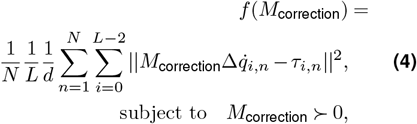

where *i* indices timesteps, *n* indices gait clip, 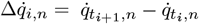 is the vector of velocity difference between step *i* and step *i*+1, 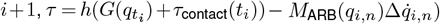 is the impulse vector that incorporates effects due to GRF, gravity and the ARB inertia. Constraining the correction inertia to be SPD ensures that the effective inertia matrix is also SPD. Next, for each gait clip *n* we can add 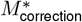 that yields *f*_min_ to the constructed *M*_ARB_(*q*_0,*n*_) to obtain the effective inertia matrix *M*_effective_(*q*_0_, *m, h*). We implemented the fitting algorithm with optimization package CVXPY (Diamond and Boyd, 2016) and CVXPYLayer (Agrawal et al., 2019).

To evaluate the physical plausibility of estimated effective inertia, we diagonally lump them (Voet et al., 2023), and for each joint aggregate relevant dimensions to compute the joint-space magnitude of inertia for that joint. We expect to see the estimated inertia matrices registering larger magnitudes on distal joints than ARB inertia’s, since we expect that our data-driven models can capture inertial coupling effects that the ARB inertia model fails to account for (Pai, 2010).

### D. The Articulated Rigid Body (ARB) Inertia Model

To compare with effective inertia, We analytically construct an ARB inertia model as a baseline, following the approach in Zell et al. (2020). First we compute the rotational inertia of each body segment *I*_torso_, *I*_pelvis_, *I*_thigh_, *I*_shank_, *I*_foot_ in its local frame under the following assumptions: (i) the pelvis and torso are elliptical cylinders; (ii) thighs and shanks are cylindrical cylinders; (iii) feet are semi-ellipsoids. This gives us the half-body rotational inertia 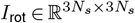, where *N*_*s*_ is the number of segments in our ARB skeletal model. Then we put translational and rotational inertia together into the spatial inertia matrix

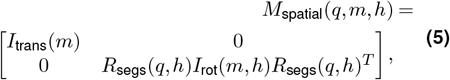

where 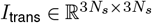 is the diagonal matrix of per-segment masses

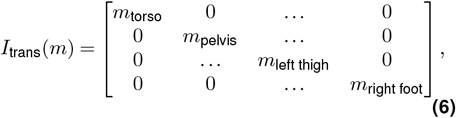

and *R*_segs_(*q*) is the 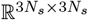 segment rotation matrix that depends on configuration *q*. The mass of each body segment is computed by multiplying total body mass *m* with Dempster’s body parameters (Winter, 2009), which is estimated from general human population. We can then compute a joint-space, pose and body proportion dependent ARB inertia matrix *M*_ARB_(*q, m, h*) for any pose *q*

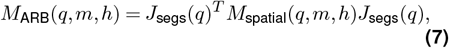

where *J*_segs_(*q*) is the 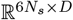 segment Jacobian matrix.

### E. Simulation

To reconstruct a subject’s motion, we inject the estimated inertia matrix *M*_effective_(*q*_*t*=0_) to a forward dynamics pipeline, and obtain pose *q* over *L* frames in the estimation window. Specifically, at each frame *t* we solve for generalized acceleration first, using Tikhonov-regularized least squares since the system can be highly ill-conditioned

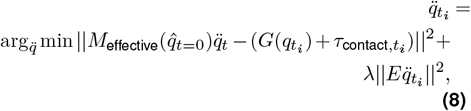

where *λ* = 1*e* − 5 is the regularization parameter and *E* is the *D* × *D* identity matrix that penalizes large joint acceleration along any particular dimension. We then apply Semi-Implicit Euler integration to update the generalized state.

### F. Metrics

#### Impulse root mean squared error

Since no ground truth inertia matrices that capture non-rigidness and inertial coupling are available, to quantitatively evaluate how close our data-driven inertia models fit to data, we resort to evaluating the closeness of predicted impulse to ground truth impulse (generalized force multiplied with time) with root mean squared error (RMSE)

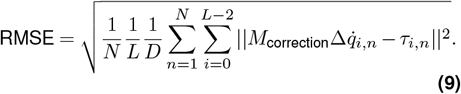

Since our generalized coordinates *q* and its derivative 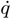 are *D*-dimensional vectors, we can break down the metric to measure root mean squared error along each degree of freedom

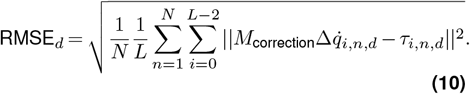

#### Mean angle error

We inject inertia matrices into a motion simulation pipeline to reconstruct lower-body kinematics, and measure the difference between predicted and ground-truth joint angles, with mean angle error (MAE), which had been established as the predominant metric for evaluating the quality of human motion prediction (Lyu et al., 2022). For a batch of *N* gait clips, we compute MAE as

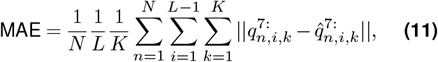

where *i* is the frame index and *k* is the joint index. Since the first six degrees of freedom correspond to root translation and orientation, we extract the later 18 DoF for measuring MAE.

## Results

### A. Joint-space Inertia

Fig. 2 shows the distribution of joint-space effective inertia estimated from constant inertia models along with inertia computed by the ARB inertia model, for each kinematic degree of freedom. Each distribution is calculated from inertia of 13 subsets of gaits; each subset consists of samples from one test subject, with either left or right stance leg. To visualize a *D* × *D* joint-space inertia matrix, we first diagonally lump entries along each row (Voet et al., 2023), then for each joint, aggregate inertia of angles that belong to the joint with 2-norm. Looking at the mean effective inertia of distal joints, i.e. left and right ankle, we see that estimated inertia are in general larger than their counterparts constructed under the ARB assumptions. This observation is consistent with what we can read from Tab. 1: (a) for walking motion, left ankle inertia is 15x larger when estimated (10.7 *kg* · *m*^2^) than constructed (0.7 *kg* · *m*^2^), and right ankle inertia is 16*x* larger when estimated (9.4 *kg* · *m*^2^) than constructed (0.6 *kg* · *m*^2^); (b) for running motion, left ankle inertia is 52x larger when estimated (43.5 *kg* · *m*^2^) than constructed (0.7 *kg* · *m*^2^), and right ankle inertia is 70x larger when estimated (42.2 *kg* · *m*^2^) than constructed (0.6 *kg* · *m*^2^). Tab. 1 shows that while inertia generally increases across all stance leg joints from ARB inertia model to data-driven model, ankle inertia increases more than mean kee or hip inertia: e.g. for the Running Set, the ARB model’s mean ankle inertia is only 0.4% of mean root translational inertia, but the data-driven model’s mean ankle inertia is 14% of mean root translational inertia.

**Table 1:** Distribution of joint-space inertia’s magnitudes, grouped by root translation, rotation and joints, of ARB inertia model vs data-driven inertia model. (a) Walking inertia magnitudes; (b) Running inertia magnitudes. Each distribution was calculated from inertia estimates over 26 subsets (13 subjects, left and right stance subset); we show mean, standard deviation of each inertia model’s estimates, and the *p*-values assuming the data-driven model yields the same mean as the ARB inertia model.

| Dimension | ARB Inertia | | Effective Inertia | | $p$ |
| --- | --- | --- | --- | --- | --- |
|  | Mean | SD | Mean | SD |  |
| (a) Walking Set |  |  |  |  |  |
| Root Trans. | 125.7 | 14.6 | 202.4 | 59.7 | 0.000 |
| Root Rot. | 79.0 | 21.2 | 133.6 | 66.9 | 0.001 |
| Pelvis | 39.5 | 11.4 | 54.7 | 28.2 | 0.017 |
| Left Hip | 27.7 | 7.6 | 50.5 | 22.0 | 0.000 |
| Left Knee | 9.3 | 2.6 | 19.7 | 11.7 | 0.000 |
| Left Ankle | 0.7 | 0.2 | 10.7 | 9.1 | 0.000 |
| Right Hip | 26.4 | 7.7 | 67.0 | 59.7 | 0.002 |
| Right Knee | 8.7 | 2.7 | 38.6 | 32.9 | 0.000 |
| Right Ankle | 0.6 | 0.2 | 9.4 | 9.6 | 0.000 |
| (b) Running Set |  |  |  |  |  |
| Root Trans. | 125.2 | 14.4 | 296.6 | 118.5 | 0.000 |
| Root Rot. | 83.8 | 21.3 | 201.4 | 93.5 | 0.000 |
| Pelvis | 42.7 | 10.0 | 89.8 | 48.1 | 0.000 |
| Left Hip | 29.3 | 8.5 | 92.7 | 60.3 | 0.000 |
| Left Knee | 10.1 | 3.0 | 64.3 | 51.6 | 0.000 |
| Left Ankle | 0.7 | 0.2 | 43.5 | 29.3 | 0.000 |
| Right Hip | 28.1 | 8.4 | 111.7 | 65.4 | 0.000 |
| Right Knee | 9.4 | 2.6 | 80.9 | 57.6 | 0.000 |
| Right Ankle | 0.6 | 0.2 | 42.2 | 29.6 | 0.000 |

**Figure 2:**
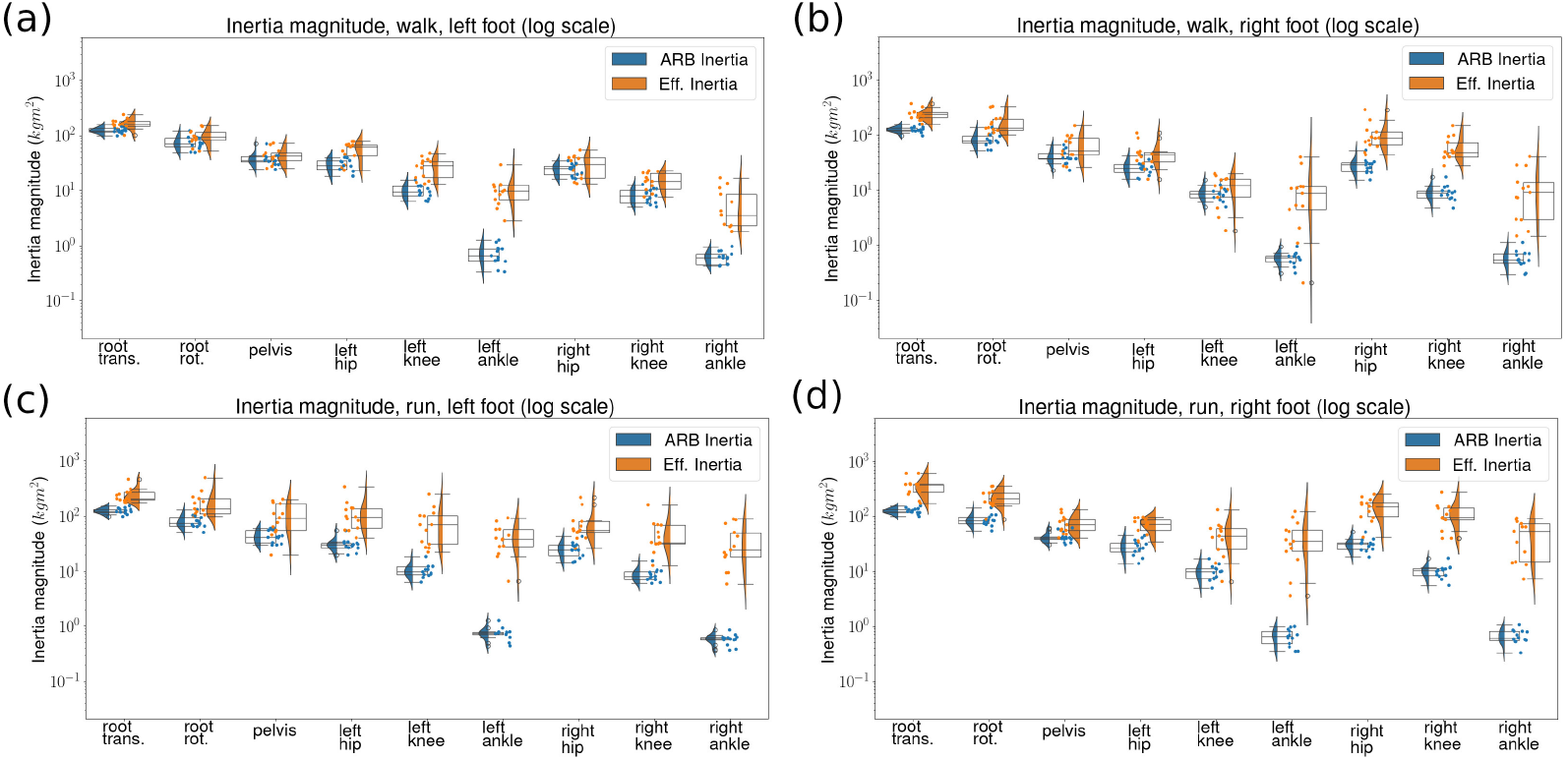
Distribution of inertia along each kinematic DoF, across subsets, where each subset contains gait samples from one subject and one stance leg. We plot inertia estimated by the constant inertia model side-by-side with those computed by the ARB inertia model. Each subject is (a) walking with left foot on ground; (b) walking with right foot on ground; (c) running with left foot on ground; (d) running with right foot on ground. The magnitudes are computed by diagonally lumping inertia matrices first, then aggregating over all angles (displacements) associated with a joint (direction). We show raw data with scattered points and distribution with box plots and violin plots. Magnitudes of inertia are shown in log scale.

From Fig. 2 we see that our estimated effective inertia is bilaterally symmetric: joints on stance legs (hip, knee and ankle) register slightly larger inertia than their counterparts on swing legs. This bilateral symmetry is not apparent in inertia constructed with the ARB model. Fig. 3 shows that the magnitude of effective inertia is larger in running than in walking, which is not the case for the ARB inertia. This is consistent with our observation from Tab. 2: in (a) running inertia are larger along all dimensions (*p <* 0.05) for the data-driven model, but in (b) the *p* values are not small enough for us to say the same for the ARB inertia model.

**Table 2:**
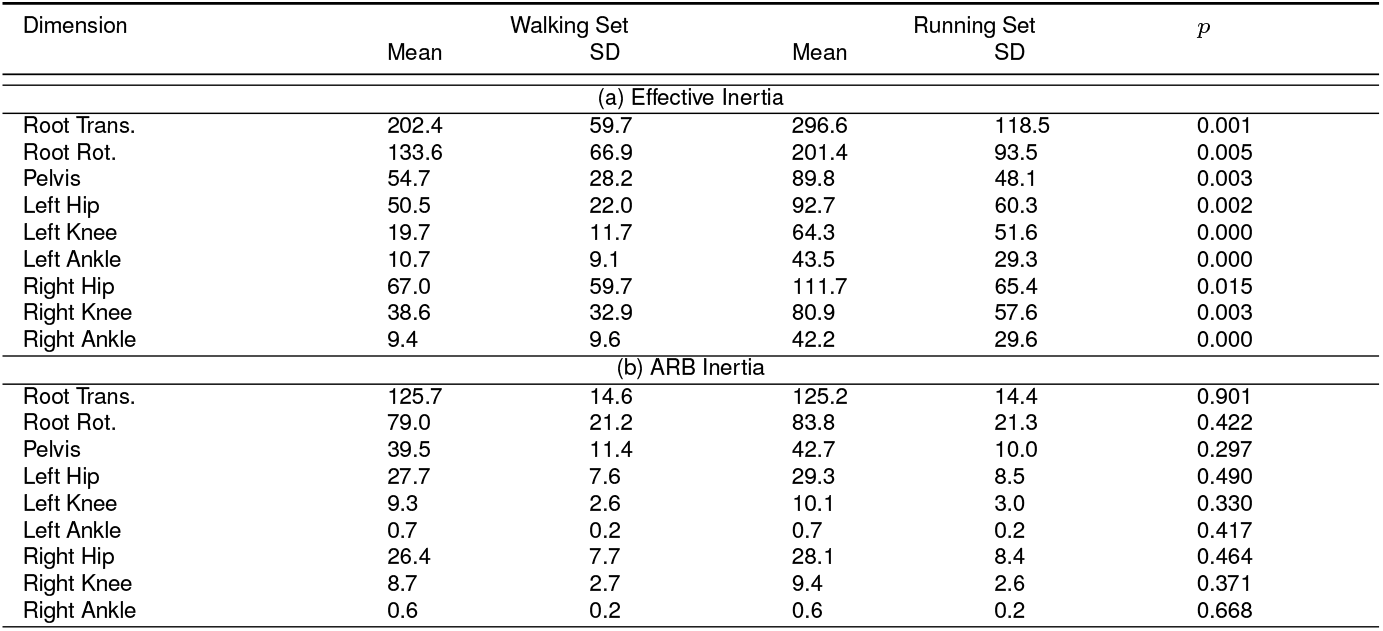
Distribution of joint-space inertia’s magnitudes, grouped by root translation, rotation and joints, of walking inertia vs running inertia. (a) Estimates from the data-driven model; (b) Constructions by the ARB inertia model. Each distribution was calculated from inertia estimates over 26 subsets (13 subjects, left and right stance subset); we show mean, standard deviation of inertia estimates for each type of motion, and the *p*-values assuming running inertia shares the same mean as walking inertia.

**Figure 3:**
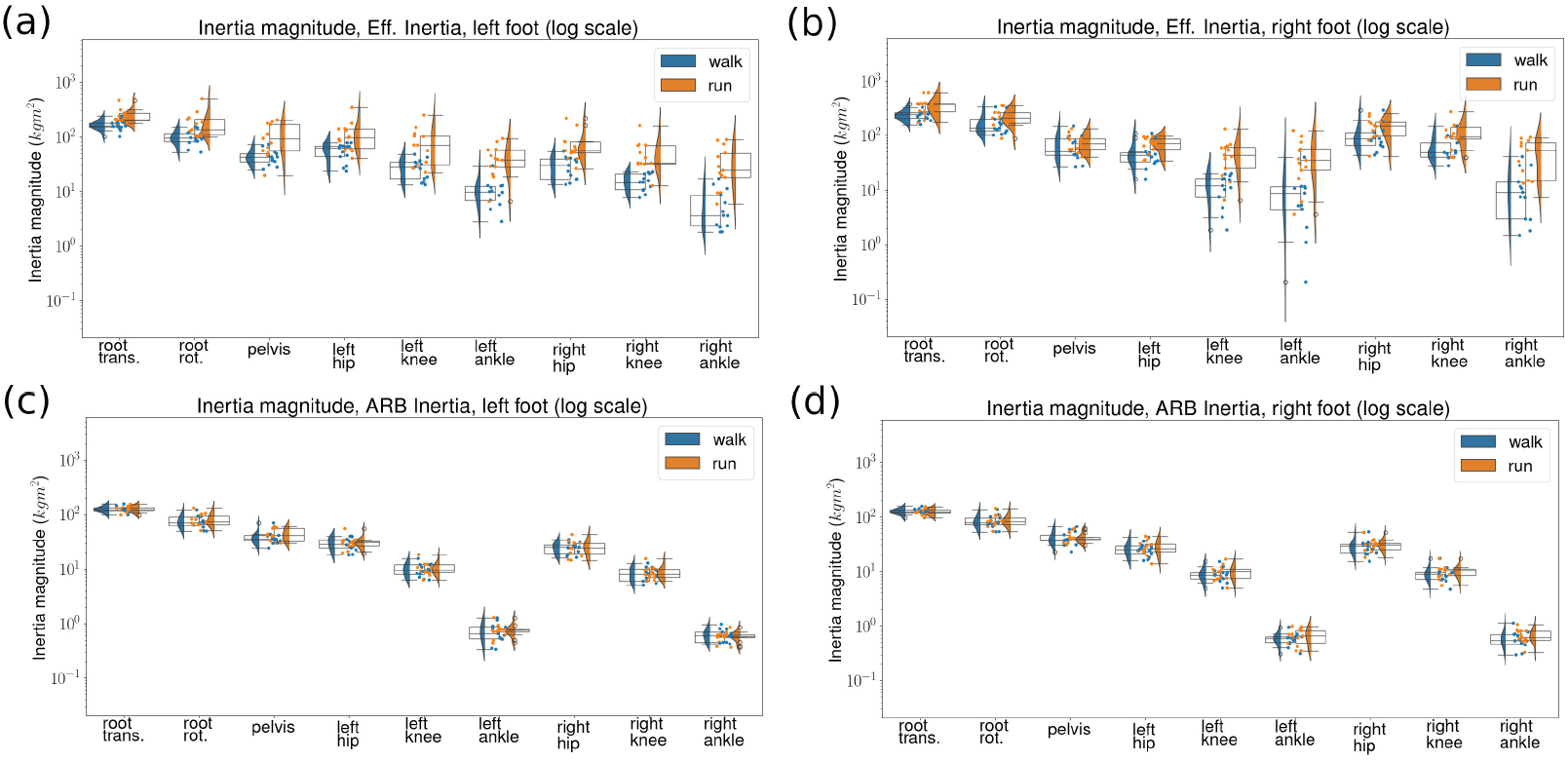
Distribution of inertia along each kinematic DoF, across subsets, where each subset contains gait samples from one subject and one stance leg. For each inertia model, We plot walking inertia side-by-side with running inertia. Data-driven models’ estimation on subsets with (a) left stance leg and (b) right stance leg; ARB inertia model’s estimation on subsets with (c) left stance leg and (d) right stance leg. The magnitudes are computed by diagonally lumping inertia matrices first, then aggregating over all angles (displacements) associated with a joint (direction). We show raw data with scattered points and distribution with box plots and violin plots. Magnitudes of inertia are shown in log scale.

### B. Quality of Impulse Fit and Kinematic Predictions

Fig. 4 shows the distribution of root-mean-squared error between predicted (constructed) impulse and ground truth impulse across 26 subsets (13 subjects, left and right stance leg subset for each), for both the data-driven inertia model and the ARB inertia model, per degree of freedom. Fig. 5 shows the distribution of MAE for motion reconstructed with the data-driven inertia model and with the ARB inertia model. Our data-driven models yield lower RMSE on the majority of degrees of freedom (*p <* 0.05): Root translation in *x, y* direction, root rotation on the the Frontal plane, pelvis rotation on the Frontal plane, hip rotation on all three planes, and knee rotation on the Frontal plane. RMSE of root translation in *z* direction, root rotation on the Sagittal plane, pevlis, knee and ankle rotation on the Sagittal plane are comparable for the two models. Looking at the mean angle error per degree of freedom, we see that the data-driven models yield lower errors on all joints and all anatomical planes (*p <* 0.05), except being tied to the ARB inertia model on the pelvis and the knee joint angles on the Sagittal plane.

**Figure 4:**
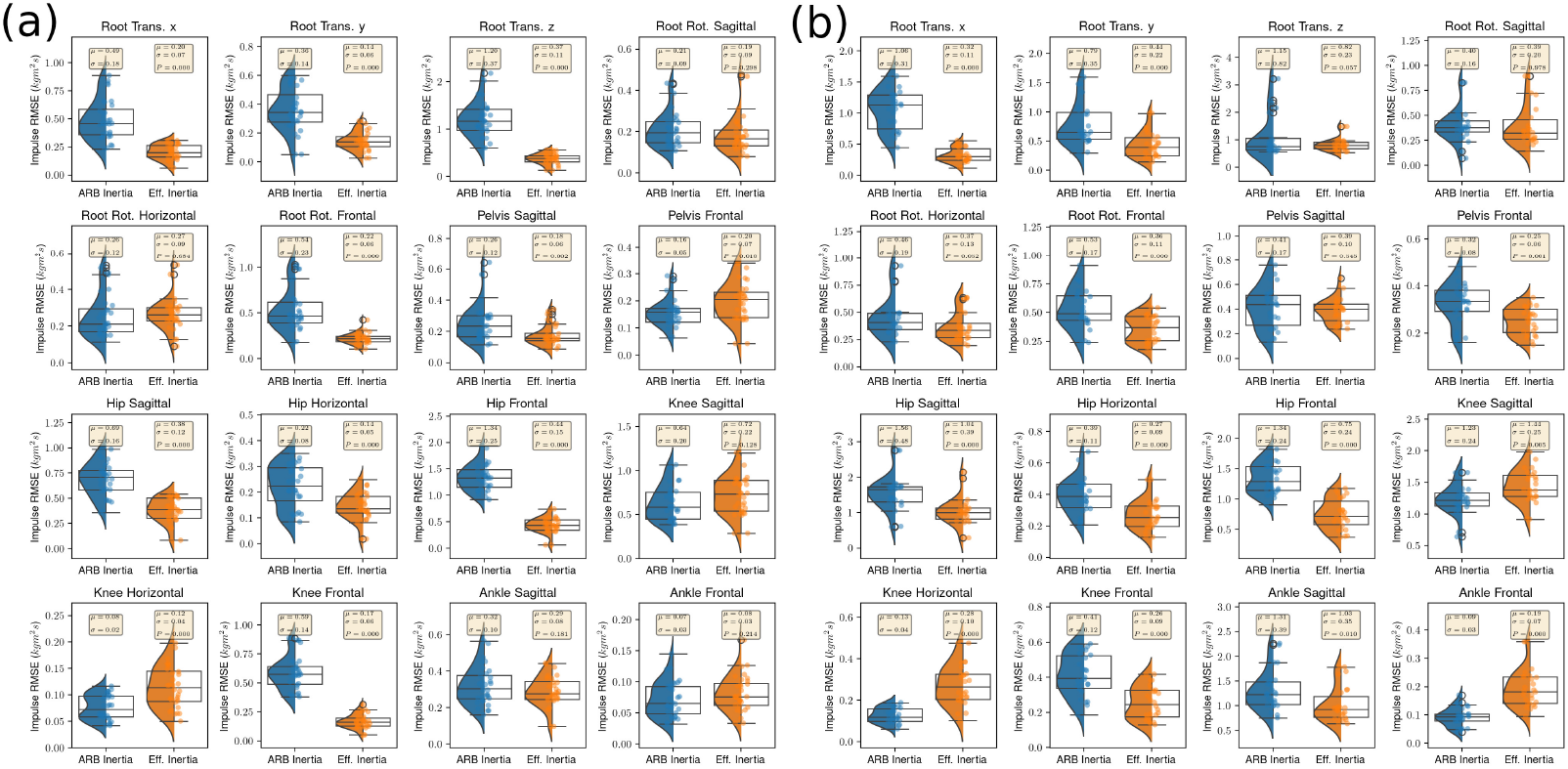
Per-DoF distribution of Impulse RMSE across training subsets. Errors from left and right leg joint angles are merged to a single dimension. (a) Walking distribution; (b) Running distribution. We show raw data points with a strip plot and distributions with box plots and violin plots. On top of each distribution with a text box we show the mean and standard deviation of RMSE across subsets; for the data-driven model we show the *p*-value assuming the data-driven model yields the same error as the ARB Inertia model. RMSE’s are plotted on linear scale.

**Figure 5:**
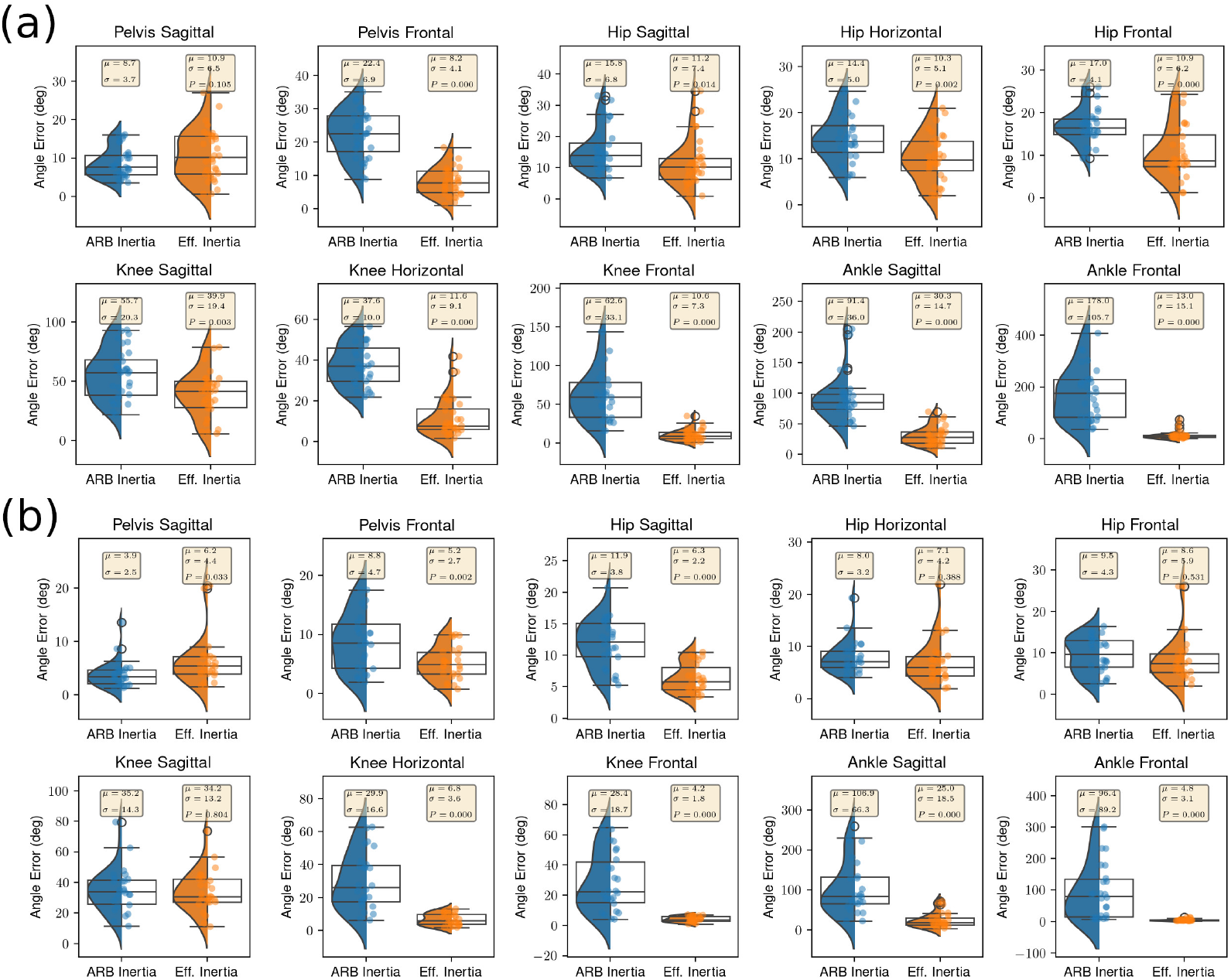
Per-DoF distribution of MAE across training subsets. Left and right leg joint angles are merged to a single dimension. (a) Walking distribution; (b) Running distribution. We show the mean, standard deviation of each distribution, and the *p*-value assuming the data-driven model yields the same error as the ARB Inertia model. MAE’s are plotted on linear scale.

Fig. 6 shows the evolution of joint angles over time for reconstructed and ground truth test set motions. The ARB inertia model diverges significantly more on knee along the Horizontal and Frontal plane, and on ankle along the Sagittal and Frontal plane. Our estimated inertia also lead to lower errors on pelvis along the Frontal plane, and on hip, knee and ankle along the Sagittal plane.

**Figure 6:**
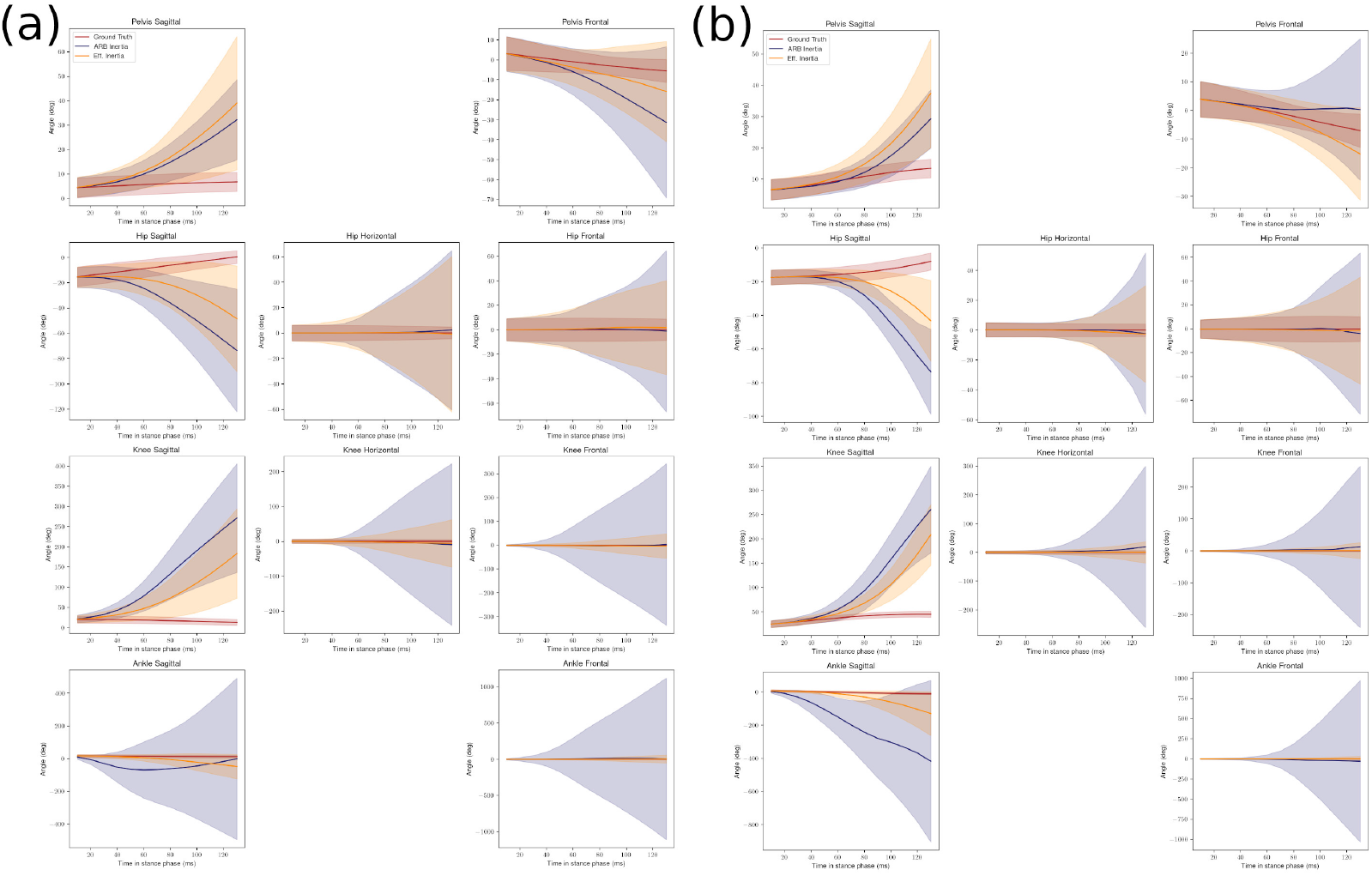
Joint angles’ mean ± one standard deviation vs. time from gait clips reconstructed by different models vs. ground truth joint angles. (a) Walking Set distribution; (b) Running Set Distribution. Distributions are calculated across gait clips from all subsets; hip, knee and ankle refer to joints on the stance leg. We initialize reconstruction at time *t* = 0 *ms* from ground-truth joint angles, so we only show differences in *t* = [10, 130] *ms*.

### C. Generalizability

Tab. 3 shows the RMSE and MAE computed over all training and test subsets, for each inertia model, broken down by degree of freedom. We notice that in terms of impulse fit, for the data-driven model RMSE increases from training set to test set, and this is more obvious on running data than on walking data. Looking at mean angle error, we see that for both the data-driven model and the ARB inertia model performance does not change much from training set to test set.

**Table 3:** Train and Test Set (a) RMSE, (b) MAE of different inertia models. For each kinematic degree of freedom, We aggregate results from all subjects and from both legs.

| Dimension | Walking Set |  | Running Set |  |
| --- | --- | --- | --- | --- |
|  | Estimated Inertia<br>Train/Test | ARB Inertia<br>Train/Test | Estimated Inertia<br>Train/Test | ARB Inertia<br>Train/Test |
| (a) RMSE ($kgm^2/s$, ↓) | | | | |
| Root Trans. x | 0.224/0.462 | 0.446/0.393 | 0.341/2.289 | 1.107/1.002 |
| Root Trans. y | 0.153/0.383 | 0.402/0.352 | 0.427/1.453 | 0.738/0.711 |
| Root Trans. z | 0.409/0.750 | 1.221/1.136 | 0.777/2.805 | 0.744/0.839 |
| Root Rot. Sagittal | 0.186/0.268 | 0.203/0.144 | 0.336/1.830 | 0.372/0.364 |
| Root Rot. Horizontal | 0.266/0.444 | 0.224/0.249 | 0.330/1.760 | 0.395/0.365 |
| Root Rot. Frontal | 0.216/0.382 | 0.463/0.434 | 0.379/1.508 | 0.541/0.539 |
| Pelvis Sagittal | 0.158/0.251 | 0.240/0.197 | 0.390/1.694 | 0.431/0.398 |
| Pelvis Frontal | 0.210/0.217 | 0.142/0.154 | 0.257/1.132 | 0.335/0.370 |
| Hip Sagittal | 0.399/0.484 | 0.709/0.683 | 0.997/2.076 | 1.500/1.514 |
| Hip Horizontal | 0.156/0.177 | 0.237/0.198 | 0.267/0.916 | 0.398/0.363 |
| Hip Frontal | 0.478/0.572 | 1.357/1.251 | 0.790/1.583 | 1.365/1.278 |
| Knee Sagittal | 0.772/0.779 | 0.657/0.580 | 1.466/1.773 | 1.197/1.266 |
| Knee Horizontal | 0.119/0.173 | 0.078/0.070 | 0.270/1.232 | 0.133/0.132 |
| Knee Frontal | 0.181/0.327 | 0.574/0.546 | 0.272/1.519 | 0.427/0.419 |
| Ankle Sagittal | 0.304/0.323 | 0.346/0.278 | 0.968/1.348 | 1.270/1.192 |
| Ankle Frontal | 0.083/0.157 | 0.078/0.052 | 0.187/1.070 | 0.092/0.068 |
| (b) MAE (deg, ↓) |  |  |  |  |
| Pelvis Sagittal | 11.43/11.05 | 9.04/8.34 | 5.03/5.69 | 3.33/4.11 |
| Pelvis Frontal | 9.09/7.81 | 21.61/22.18 | 5.18/5.64 | 8.91/8.64 |
| Hip Sagittal | 9.64/10.13 | 16.40/12.06 | 6.38/6.14 | 11.93/12.12 |
| Hip Horizontal | 10.89/7.48 | 15.30/13.14 | 6.51/7.09 | 7.59/8.13 |
| Hip Frontal | 11.65/11.27 | 15.80/15.95 | 7.11/7.31 | 9.66/7.72 |
| Knee Sagittal | 42.74/39.29 | 59.44/56.27 | 32.80/34.50 | 33.48/36.54 |
| Knee Horizontal | 10.25/6.55 | 36.29/35.52 | 6.92/8.25 | 30.36/28.80 |
| Knee Frontal | 10.46/10.77 | 53.48/61.46 | 4.15/4.58 | 28.78/28.46 |
| Ankle Sagittal | 32.59/24.63 | 88.20/86.25 | 19.48/22.84 | 101.79/124.77 |
| Ankle Frontal | 8.75/9.03 | 151.21/183.15 | 4.14/5.70 | 98.66/127.61 |

## Discussion

### Inertial coupling

Fig. 2 shows estimated ankle inertia is on average 41 times higher than analytically constructed ankle inertia. Specifically, the ARB ankle inertia are minuscule and in fact nearly negligible compared to inertia of other joints, e.g. for walking gait clips with left stance leg, mean left ankle inertia is 0.72 *kgm*^2^ whereas mean pelvis inertia is 37.9 *kgm*^2^; for the data-driven model we see ankle inertia becoming considerably larger when compared with inertia along other degrees of freedom, e.g. 6.2 *kgm*^2^ of left ankle inertia compared with 13.7 *kgm*^2^ of pelvis inertia. Pai (2010) used a rather simple musculoskeletal model consisting of two joints and a monoarticular muscle to show that when muscle mass is unlumped from bone mass, the muscle increases inertia of joints distal to it. In the case of human ankle, unlumping muscle mass of triceps surae alone can contribute 7.6% of additional inertia (Pai, 2010). Our results can confirm that ankle inertia estimated from real-world data without the lumping assumption is larger than the lumped, ARB ankle inertia, and we find it reasonable to believe that the additional estimated inertia is due to shank muscles’ coupling to the ankle joint. However the precise contribution of each muscle to the increased inertia is yet to be determined and requires further study.

### Bilateral symmetry

We hypothesize that the the symmetry is due to the role of neural activation on inertia coupling. The difference in active muscle groups on the stance vs. swing leg (Bonnefoy-Mazure and Armand, 2015) may have led to different inertial coupling effects and thus, have led to larger inertia on stance leg joints. More research to test this hypothesis is still needed.

### Motion-type dependence of inertia

Another aspect the corrective inertia is able to capture is the dependence of effective inertia on motion type. Although we can easily deduce that ARB inertia model cannot capture motion type dependence because it is not fitted on data and it is not parametrized by motion type, we cannot tell much about why the estimated running inertia are larger in magnitudes, beyond hypothesizing that this is related to muscle activation patterns being different in running than in walking.

### Quality of estimation and implication of using different inertia estimates to reconstructing lower body Kinematics

Since the data-driven models yield lower or comparable RMSE for the majority of kinematic degrees of freedom (with *p <* 0.05) we can confirm that data-driven models yield more accurate estimates. Using different inertia estimates in simulation can lead to drastically different performances on reconstructing lower body kinematics: using joint-space inertia matrices estimated from data yields lower mean angle errors for both walking and running motion, regardless whether we are reconstructing training gait clips used to fit the models or testing gait clips. Moreover, gaits reconstructed with ARB inertia becomes unstable sooner than gaits reconstructed with data-driven models, as shown and discussed with Fig. 6. This is due to ARB inertia matrices being more ill-conditioned, will condition numbers order(s) of magnitude larger than effective inertia matrices. So using estimated inertia improves the accuracy and stability of lower body kinematic reconstruction.

### Generalizability

The data-driven model yields larger motion errors than the ARB models on test subsets: see Tab. 3. This is expected since (i) test gait clips were not used to fit data-driven models; (ii) our data-driven models are not parametrized, so they cannot infer inertia from test gait clips’ kinematic and dynamic information.

## Conclusion

We developed a data-driven model for representing subject-specific effective joint-space inertia matrix of the lower body of a human in everyday walking and running, and a method to fit such models to kinematic and dynamic data of human motion. The data-driven model predicts an effective joint-space inertia matrix that is the sum of an ARB inertia matrix and a correction inertia matrix which accounts for effects overlooked by ARB assumptions that approximate physical and geometric properties of human body segments. The correction inertia assumes nothing other than that it is symmetric and positive definite, yet reveals effects of inertial coupling that inertia matrices constructed under the ARB assumptions can fail to capture (Pai, 2010). With the RMSE metric we confirmed that our data-driven model can fit to data well, and with the MAE metric we have demonstrated that injecting estimated inertia into a simulation framework yields lower error in reconstructing lower body kinematics.

## ACKNOWLEDGEMENTS

We thank the Natural Sciences and Engineering Research Council of Canada (NSERC) for its funding of the project. Also, we extend our gratitude to Felix Winkler for his help with the data processing pipeline.

